# Robustness tuning: mechanisms of acclimation-driven plasticity in a central pattern generator

**DOI:** 10.64898/2026.07.07.737058

**Authors:** Sonal Kedia, Max Kenngott, J.D. Rittenberg, Eve Marder

## Abstract

Temperature influences neuronal and circuit output and extreme temperatures can disrupt neuronal performance. Acclimation invokes a form of neuronal plasticity that we call ‘robustness tuning’, that preserves nervous system performance during seasonal alterations in environmental conditions. The stomatogastric nervous system (STNS) of the American lobster, *Homarus americanus*, produces stereotyped rhythmic motor patterns that are maintained over a range of acute temperature changes, but lost under more extreme conditions. In the wild, *H. americanus* experience water temperatures from ∼2°C to 25°C during the course of a year. We acclimated lobsters to 18°C versus 4°C for ∼3 weeks, and found that the pyloric rhythm from warm-acclimated animals maintained its characteristic properties over an extended temperature range, when compared to those recorded from cold-acclimated animals. There were acclimation and temperature dependent differences in the responses of pyloric neurons to the neuropeptide, Crustacean Cardioactive Peptide (CCAP). Computational models suggest that pyloric neuron morphology and neuromodulator conductance distribution play a role in robustness tuning, the reversible changes that allow animals to repeatedly adapt to seasonal change.

## Introduction

Nervous systems are highly dynamic and are shaped and reshaped by experience throughout the animal’s lifetime. Animals learn new skills and associations that are stored using long-term plasticity mechanisms introducing permanent changes in neuronal circuitry [1–6]. Balancing the accrual of activity changes are homeostatic mechanisms that renormalize circuit activity to set points [7–17]. This taxonomy, however, fails to address circumstances in which plasticity generates persistent, yet reversible changes to network behavior, such as those seen in seasonal acclimation to temperature changes that occur cyclically through a long-lived animal’s lifetime.

All physiological processes function well within some range of environmental factors with distinct limits, beyond which they fail [18]. However, these limits are not fixed and can be adjusted throughout an animal’s lifetime to reflect long-term changes in the environment. Thermal acclimation is a process by which animals adapt to new, prolonged temperature regimes by shifting the limits of temperature at which they function [19]. Although there is an enormous literature on acclimation [20], less is understood of the circuit and cellular mechanisms by which animals achieve it. We use the term ‘robustness tuning’, loosely derived from control theory concepts, for the plasticity through a network adapts its permissive range based on environmental constraints.

The American lobster, *Homarus americanus,* lives in waters that can range from 0-25°C across seasons making it ideally suited for addressing the underlying mechanisms of neural acclimation [21]. The stomatogastric ganglion (STG) of lobsters contains a central pattern generating circuit whose output must be maintained throughout the entire life of the animal, despite changes in the physical environment [22–24]. The pyloric circuit of the STG produces nearly uninterrupted rhythmic contractions of the pylorus, a filtering structure in the foregut, by a characteristic triphasic pattern of activity of its individually identifiable constituent neurons [24–26]. The frequency of the pyloric rhythm increases monotonically with increasing temperature, before losing coherence at and above a ‘crash’ temperature [27]. The thermal limits of this network, or the crash temperature, change upon acclimation [27–30].

We acclimated animals to different temperatures and probed circuit and cellular changes underlying this flexibility through electrophysiological recordings and used computational models to suggest possible mechanisms for these changes.

## Results

### Hot acclimation extends the thermal performance range of the pyloric rhythm

Our experimental strategy was to describe thermal acclimation in the *H.americanus* STNS, and then to dissect its mechanisms at the circuit and cellular levels. The STNS was dissected from adult *H. americanus* housed at 4°C (‘cold’ acclimation) or 18°C (‘hot’ acclimation). The triphasic pyloric rhythm was recorded extracellularly from the lateral ventricular nerve (*lvn*) which carries axons of the-pyloric dilators (PD), lateral pyloric neuron (LP) and pyloric neurons (PYs) (Fig 1a, left panel) [23]. Spikes from each neuron type were identified and recorded as the entire preparation was exposed to gradually increasing saline temperatures until a ‘crash’ was observed-either the bursting slowed down or entire units disappeared. Example recordings show a preparation from a cold acclimated lobster crashed at 24°C, whereas recordings from a hot acclimated lobster showed consistent triphasic activity up to 31°C (Fig 1a).

**Figure 1.**
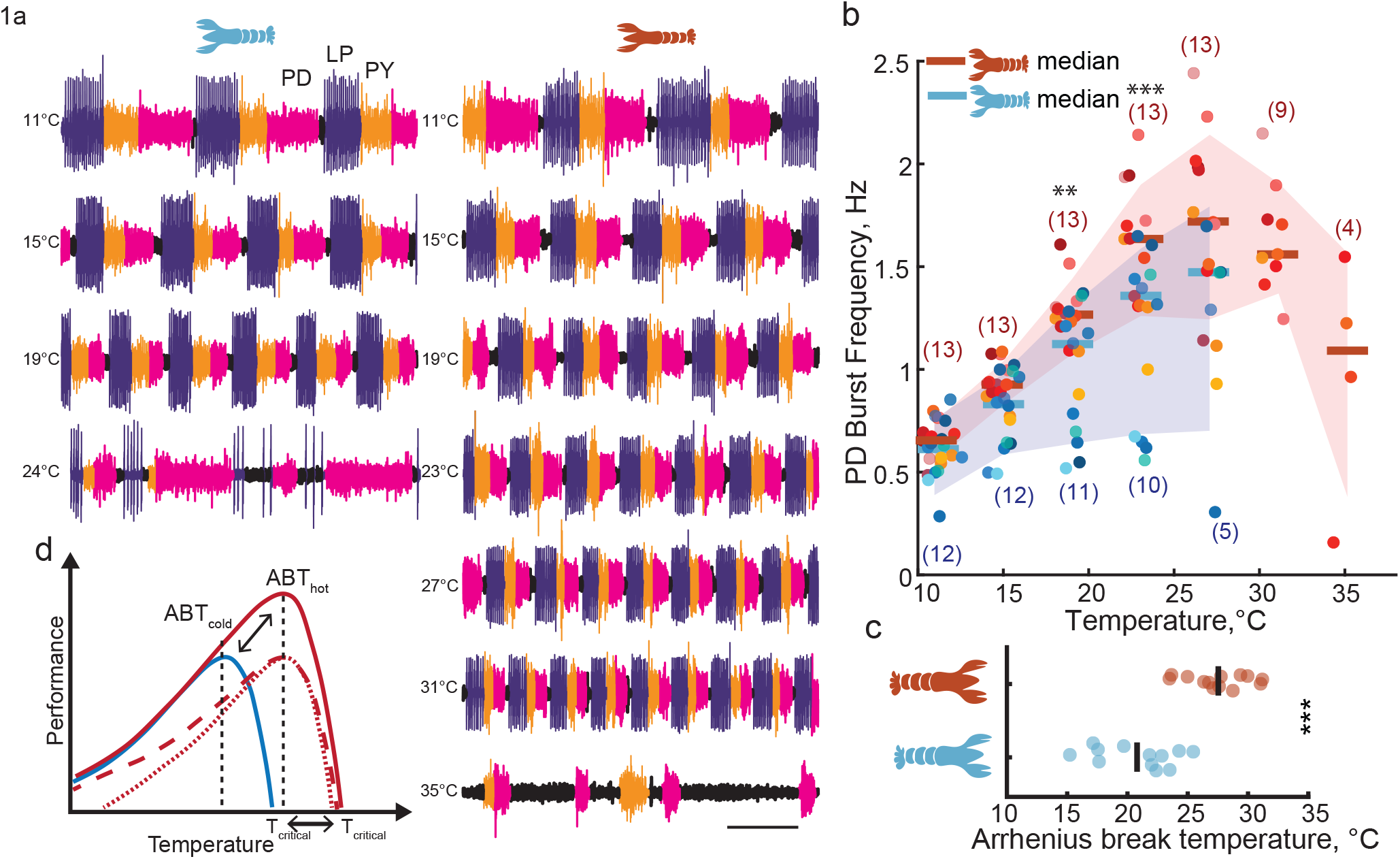
Acclimation changes thermal limits of the pyloric network. a. Sample traces of extracellular recordings from lvn from a cold animal (left) and hot animal (right) taken at increasing temperatures. Calibration: 1s. b. PD burst frequencies versus temperature across all animals in both groups. Cool colors indicate individual cold animals and warm indicate hot animal data points. Ns in parenthesis. c. Population averages (vertical lines) and individual animal distributions (circles) for ABTs in the cold (N=12) and hot groups(N=13). d. Schematic of potential patterns of shifting thermal limits after acclimation-cold acclimation represented in blue, hot in various red lines. * p<0.05, ** p<0.01, *** p<0.001

PD burst frequency was quantified in the two groups in response to temperature (Fig. 1b). Burst frequencies were compared over 11-23°C when most preparations were still active. Both groups responded to increasing temperature with an increase in burst frequency, consistent with published reports of temperature effects on the pyloric rhythm in multiple decapod species [27–32]. There was a strong effect of temperature on burst frequency from 11°C to 23°C (Table ST1). At moderate temperatures (11–15°C), pyloric rhythm frequencies did not differ between groups. Preparations from hot-acclimated animals had higher burst frequencies at temperatures ≥16°C than preparations from cold-acclimated animals (Table ST2). At higher temperatures preparations from cold-acclimated animals slowed or ‘crashed’ and were not recorded further.

The Arrhenius Break Temperature (ABT) is defined as the point in the Thermal Performance Curves (TPC) (Fig. 1b) where frequencies no longer obey the Arrhenius equation, that is, they slow instead of increasing frequency with temperature increases (Methods). The ABT is a useful quantification of the limits of optimal performance for a system. It is less subjective than ‘crashes’ and Q_10_s are a valid measurement only up to this temperature. The ABT of preparations from cold acclimated animals were significantly lower than the ABT of preparations from hot acclimated animals (Student’s T-test, p=<0.0001) (Fig. 1c).

Thermal performance curves can have different patterns of changes post acclimation to accommodate an increase in thermal limits (dotted/dashed red lines, Fig. 1d); each of these patterns suggests different molecular mechanisms [33]. The response of the pyloric network to acute temperature changes is largely unaltered by acclimation (Q_10_s are not significantly different between ‘hot’ and ‘cold’ groups between 11-21°C), but the operational range for producing a pyloric rhythm is enlarged (solid red line, Fig 1d).

### Network output and dose responses to CCAP are affected by temperature and acclimation

The above experiments confirmed that animals acclimated to different temperatures exhibit different degrees of thermal resilience. Compared to preparations drawn from animals in the cold acclimation group, preparations from the hot acclimated animals had a higher ABT, and correspondingly an extended permissive thermal range.

We next investigated the circuit level mechanisms of this range acclimation, specifically the role of neuromodulation. Neuromodulation exhibits strong seasonal trends in eurythermal ectotherms that helps coordinate the timing of behavioral programs influenced by temperature [34–36]. Neuromodulation plays an important role in generating the pyloric rhythm [37–40] and in extending the temperature range over which the circuit is operational [41, 42]. We hypothesized that crashes of the pyloric rhythm at high temperatures after cold acclimation could arise from acclimation-produced differences in neuromodulation. A deficiency in neuromodulator production and/or release could underlie a smaller operational range after cold acclimation. If so, adequate neuromodulator applied exogenously could restore activity at higher temperatures after cold acclimation and produce activity similar to preparations from hot-acclimated animals.

An exploration of the involvement of neuromodulation in robustness tuning was conducted through dose response curves to CCAP in preparations in which intrinsic synaptic neuromodulation was removed (decentralization). Hormonal sources of neuromodulators were already removed from STNS preparations by the dissection and decentralization further removes endogenous synaptic sources. This allows us to test the effects of controlled amounts of exogenously applied modulator to study its effects in isolation. CCAP is one of many modulators that can improve or generate a triphasic rhythm in decentralized conditions [43] and has important physiological associations with controlling cardiac frequency and molting [44, 45].

Many neuropeptides in the crab STG, including CCAP, appear to converge on a single voltage-dependent inward current, termed I_MI_ [40, 46] and can restore/maintain triphasic activity through its activation. This current was first described using the neuropeptide proctolin [47], and elements of its intracellular signaling have been characterized [48, 49]. Increasing concentrations of CCAP would theoretically activate a larger I_MI_ current, similar to the action of multiple other modulators.

### Determining dose response curves

At 10°C, circuit output was similar between the two acclimation groups and at 20°C we started to see circuit slowing and failure in the cold-acclimated group (Fig. 1b). We studied CCAP dose responses of the pyloric network at these two temperatures in both acclimation groups to dissect cold-acclimated network failure and characterized the following:

a. Changes in dose responses due to acclimation at a specific temperature
b. Changes in dose responses due to acute temperature changes
c. The interaction of acclimation and acute temperature on CCAP dose responses

Pyloric activity was recorded in decentralized preparations from cold-acclimated and hot-acclimated animals at either 10°C or 20°C (start temperatures alternated between experiments), and the preparations were exposed to increasing concentrations of CCAP, followed by a wash-out of modulator. Subsequent to a change in temperature, the dose response measurements were repeated (Fig. 2a-d: example extracellular recordings from pyloric dilator nerve (pdn) (PD spikes in magenta) and lateral pyloric-pyloric constrictor nerve (lp-pyn) (LP spikes in purple and PY spikes in orange).

**Figure 2.**
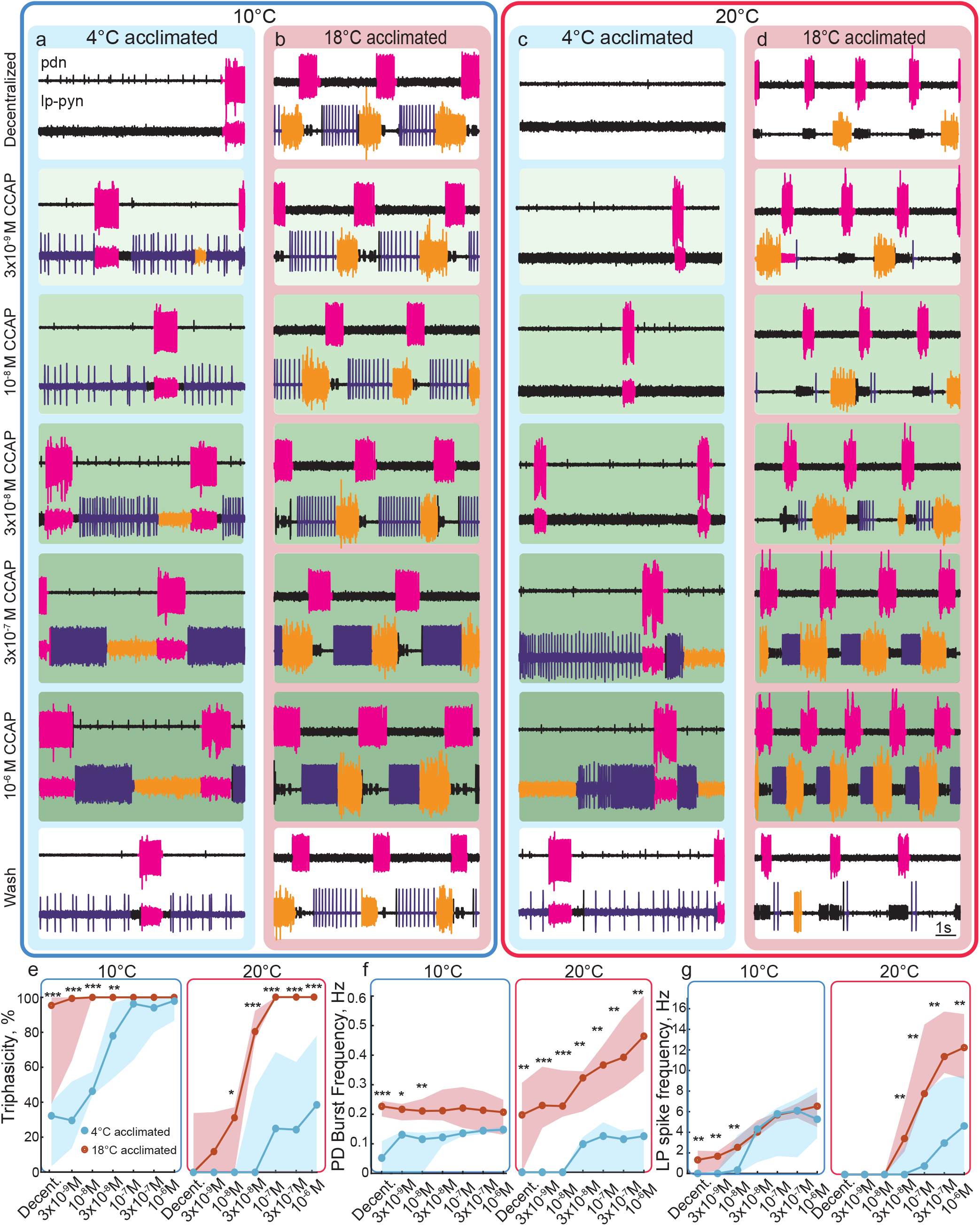
Neuromodulation has different outcomes in the two acclimation groups at different temperatures. a-d. Sample traces of extracellular recordings made from *pdn* and *lp-pyn*. Filled blue boxes contain recordings from a cold acclimated animal at 10°C (a) and 20°C (c) and the red boxes contain recordings from a ‘hot’ acclimated animal at 10°C (b) and 20°C (d). e. Group medians for triphasic transitions % at 10°C and 20°C. Red circles are hot acclimated medians; red shading is the interquartile range. In blue are values for cold animals for e-g; N=6 for each group f. Median pyloric burst frequency values in both groups at 10° and 20°C. g. Median LP spike frequencies in both groups at 10°C and 20°C. * p<0.05, ** p<0.01, *** p<0.001

In a decentralized STG from a cold acclimated animal studied at 10°C, the individual neurons were silent (LP, PY) or bursting (PD) and the network activity was not triphasic (Fig 2a, top panel). Increasing concentrations of CCAP increased bursting activity in all the pyloric neurons and restored triphasic bursting in the presence of CCAP >3×10^−8^M (Fig 2a, lower panels). At 20°C the same preparation showed reduced activity in the absence of neuromodulation (Fig 2c, top panel). Unlike at 10°C however, CCAP restored firing in all the neurons but did not restore regular triphasic activity even at the highest dose tested (10^−6^M) (Fig 2c lower panels).

In contrast, example recordings from a decentralized STG from a hot acclimated animal studied at 10°C had intact triphasic activity in the absence of neuromodulation (Fig 2b, top panel). All the neurons responded to CCAP by increasing spiking activity as in the cold-acclimated preparation, but the overall rhythm was qualitatively similar across the dose response conditions (Fig 2b, lower panels). In the same preparation triphasic activity was lost at 20°C, although PD continued to burst in the absence of CCAP (Fig 2d, top panel). At CCAP concentrations >3×10^−8^M a regular triphasic rhythm was restored (Fig 2d, lower panel).

The dose responses of 3 different parameters of circuit activity were quantified. We measured: I. the number of triphasic transitions, or triphasicity % (Methods). II. PD burst frequency III. LP neuron activity. Results are presented below broken down by test temperature and acclimation group.

### Triphasicity

The % triphasicity of the pyloric network over a 2-minute period was quantified at each dose of CCAP, at both temperatures, in both acclimation groups (Fig 2e).

#### Dose responses at 10°C

In the cold-acclimated group triphasicity was reduced to 30% when preparations were decentralized. CCAP doses (>10^−8^M) increased triphasicity which saturated at 100% by 10^−7^M CCAP (Fig 2e, left panel).

In the hot-acclimated group decentralization caused a negligible reduction in rhythmicity. All preparations were fully triphasic in CCAP doses >10^−8^M, the response was blunted due to ceiling effects (Fig 2e, left panel).

The cold acclimated group had significantly lower triphasicity than the hot when decentralized and in CCAP concentrations <10^−7^M (Table ST3). CCAP responses were larger in the cold-acclimated group and smaller in the hot such that at higher doses the two groups had similar outputs and were fully rhythmic (Fig 2e, left panel).

#### Dose responses at 20°C

As at 10°C in the cold-acclimated group triphasicity was reduced to ∼0% in decentralized preparations by raising the temperature (Fig 2e, right panel). CCAP (>10^−7^M) restored a percentage of triphasic transitions but the highest dose of CCAP (10^−6^M) tested was insufficient at restoring full rhythmicity at this temperature.

Unlike at 10°C in the hot-acclimated group triphasicity was similarly ∼0% in decentralized preparations. Preparations recovered triphasic transitions in doses as low as 3×10^−9^M CCAP and were fully triphasic at all concentrations >10^−7^M CCAP (Fig 2e, right panel).

Both acclimation groups lost rhythmicity completely at 20°C. CCAP restored triphasic transitions and the hot acclimation group had higher sensitivity and output. The hot acclimated group was significantly more triphasic at all doses >10^−8^M (Table ST3)(Fig 2e, right panel).

### PD burst frequencies

#### Dose responses at 10°C

In the cold-acclimated group PD bursting and burst frequencies were strongly reduced after decentralization (0.06± 0.05 vs ∼0.6Hz in intact networks (Table ST1)). This effect was rescued by CCAP, with PD burst frequencies increasing in all concentrations of CCAP (Fig 2f left panel, Table ST4).

In contrast, in the hot-acclimated group PD neurons continued to burst but slower in decentralized conditions than in intact (0.2± 0.1 vs ∼0.6Hz in intact networks (Table ST1)). Unlike the cold-acclimated group, PD burst frequencies were insensitive to CCAP doses (Fig 2f left panel, Table ST4).

PD burst frequencies were significantly higher in the hot acclimation group at CCAP concentrations <10^−8^M (Table ST4) and similar between groups at higher doses.

#### Dose responses at 20°C

In the cold-acclimated group PD bursts completely disappeared in decentralized conditions. PD bursts re-emerged in CCAP doses >3×10^−8^M and increased at higher doses (Fig 2f, Table ST4, 5). Despite this, at 10^−6^M CCAP PD burst frequency at 20°C remained lower than the frequencies at 10°C in the same acclimation group.

In contrast, in the hot-acclimated group PD neurons continued to burst and burst frequencies remained unchanged from 10°C to 20°C (Fig 2f, Table ST4, 5). PD burst frequencies increased in CCAP >3×10^−8^M. At 10^−6^M CCAP, PD burst frequencies at 20°C were approximately double the frequency at 10°C (Q_10_≈2) in the same acclimation group.

Burst frequencies were significantly higher in the hot-acclimated group in all conditions while sensitivities were unaltered by acclimation (Table ST4, 5) at 20°C.

### LP activity

LP activity can be tonically firing after decentralization. Therefore, we measured the total number of LP spikes/second as a measure of LP activity.

#### Dose responses at 10°C

In the cold-acclimated group the LP neuron failed to spike in decentralized conditions. Spiking increased in CCAP doses >3×10^−8^M (Fig 2g left panel, Table ST6,7).

In contrast, in the hot-acclimated group the LP neurons continued to spike after decentralization. LP spike frequencies also increased in CCAP (Fig 2g left panel, Table ST6,7).

LP spike frequencies were significantly higher in the hot acclimation group in CCAP concentrations <3×10^−8^M (Table ST7) and similar between groups at higher doses.

#### Dose responses at 20°C

In the cold-acclimated group the LP neuron continued to be silent in decentralized conditions. LP spike frequencies increased in CCAP >10^−7^M reaching the highest value of 6.5±3.7Hz at 10^−6^M CCAP (Fig 2g, left panel, Table ST7).

In the hot-acclimated group LP spikes disappeared from 10°C to 20°C in decentralized conditions (Fig 2g, left panel). LP spiking increased at lower doses of CCAP (>3×10^−8^M) and more steeply (13±3.5Hz in 10^−6^M CCAP, Table ST7).

LP spike frequencies were significantly higher in the hot-acclimated group than the cold-acclimated group at all doses of CCAP > 3×10^−8^M (Table ST7).

In conclusion, there were significant interactions between the effects of acclimation, temperature and CCAP on all features of activity. The above results support the hypothesis that modulation plays a role at the circuit level in setting the thermal limits of robust activity in acclimated animals. However, this cannot be the whole story. Although modulation expands the range at which both hot and cold acclimated preparations exhibit triphasic rhythms and PD bursting, exogenous modulation is not sufficient to turn a cold-acclimated preparation into a hot acclimated preparation. While the two groups converge in behavior at 10°C with applied CCAP, at 20°C, their the gap in performance cannot be rescued with exogenous modulation.

### LP excitability was altered by elevated temperatures, CCAP and acclimation

Based on the dose response experiments at 20°C, it appears that the effects of elevated temperature have similar effects on some aspects of network output in both acclimation groups, i.e., both triphasic activity and LP spikes were lost. Nonetheless, the two groups responded to CCAP differently at higher temperatures. This suggests that acclimation is not simply a function of neuromodulator availability.

Activity at the saturating end of the dose response may be greater after hot acclimation for numerous reasons. The direct impact of CCAP (the I_MI_ current) might be selectively increased at high temperatures after hot acclimation through changes in the neuropeptide receptor-I_MI_ pathway. Intrinsic conductance values could be different, producing higher excitability. These changes could be cryptic in baseline conditions but still interact non-linearly with the similar I_MI_ values in different acclimation groups with different resulting activities. Finally, both of these could be true.

We explored these possibilities by measuring changes in LP excitability in response to CCAP and acute temperature in the two acclimation groups. We studied spike thresholds and firing rates by injecting slow current ramps into synaptically isolated LP neurons (Methods). Intracellular LP neuron recordings from a cold-acclimated animal had similar spike threshold values at 10°C with or without CCAP (Fig 3a, left panel). Spike thresholds hyperpolarized by ∼20mV at 20°C. CCAP depolarized thresholds back to 10°C values and increased spike frequencies (Fig 3a, right panel).

**Figure 3.**
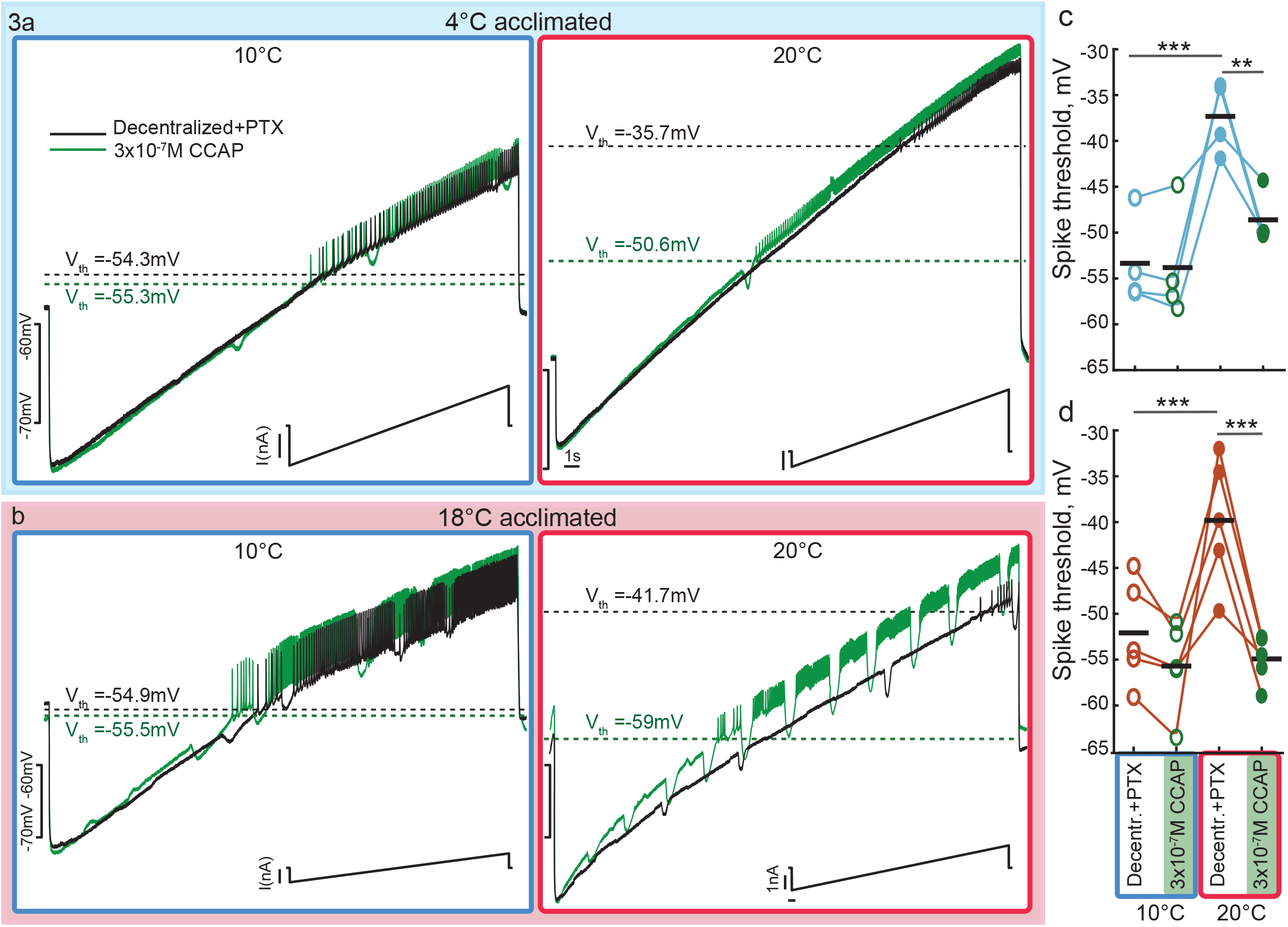
LP spike thresholds lowered by elevated temperature and recovered by CCAP in both acclimation groups. a. Sample traces from a cold-acclimated animal. Membrane voltage traces from an LP neuron (top traces) during slow ramps of current injection (bottom trace) repeated at 10°C (left panel) and 20°C (right panel) before and after applying CCAP. Spike thresholds marked with dotted lines. b. Sample traces of an LP neuron from a hot-acclimated animal with ramps as in a. c-d. Spike thresholds in cold and hot-acclimated animals respectively. Horizontal bars represent population means. Individual circles connected by lines represent one animal. ** p<0.01, ***p<0.001.

The LP neuron from a hot-acclimated animal responded to CCAP with an increase in spike frequencies at 10°C and 20°C (Fig 3b). The spike threshold was unaffected by CCAP at 10°C; at 20°C the spike threshold depolarized and CCAP application hyperpolarized the threshold (Fig 3b, right panel).

LP spike thresholds were sensitive to temperature and CCAP in both acclimation groups (Fig 3c, Table ST8, 9). LP neuron spike thresholds in decentralized conditions were significantly depolarized at 20°C relative to 10°C (p < 0.0001) in preparations from both acclimation groups (if preparations failed to spike with ramps at different rates, spike thresholds were measured by injecting short current steps of increasing values until a spike was elicited, Fig S1). In both acclimation groups, CCAP significantly hyperpolarized the spike threshold at 20°C (p = 0.0017 cold, p < 0.0001 hot) but had no effect at 10°C (Fig 3c, Table ST8). Thus, elevated temperature depolarized spike thresholds and CCAP had a similar depolarizing effect in both groups.

LP spike frequencies as a function of membrane potential were also measured during the ramps. At 10°C, LP spike frequency curves were similar with and without CCAP in the cold acclimated group. The LP spike frequency curve shifted towards higher frequencies during CCAP application in the hot-acclimated group (Fig 4a, b left panels; Table ST9, 10). These changes were quantified for membrane potentials between −50 to −45mV (Fig 4c) and compared using paired T-tests. Baseline spike frequencies were the same in the two acclimation groups. CCAP increased firing frequency in the −50 to −40mV window in cold-acclimated animals (Δ=2.2Hz, p = 0.04) and more strongly in hot-acclimated animals (Δ=8.3Hz, p = 0.012) such that in CCAP, LP spike frequencies were higher in the hot-acclimation group than in the cold (p = 0.04).

**Figure 4.**
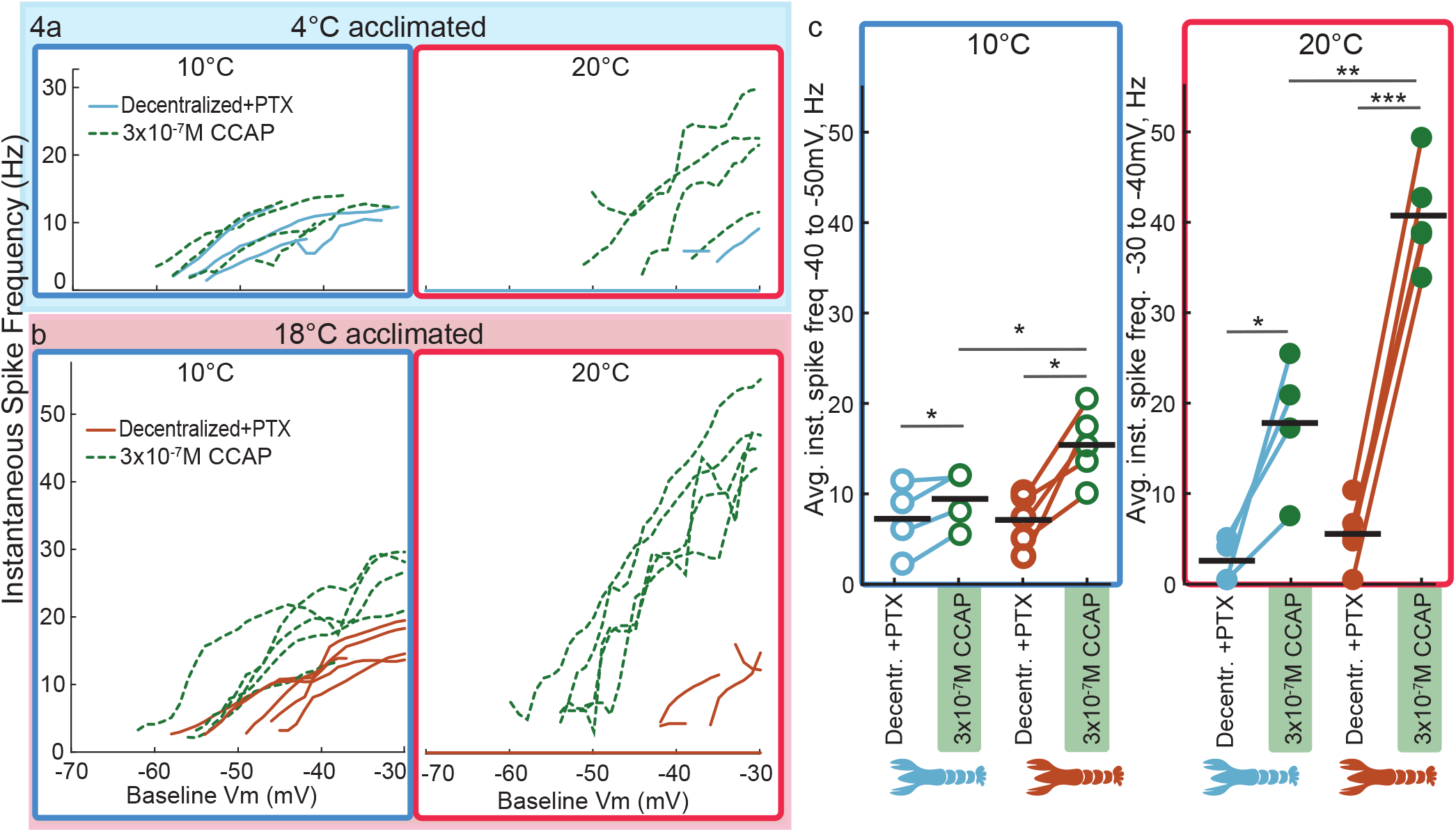
LP firing frequency is more responsive to CCAP in hot-acclimated animals at 10°C and 20°C a-b. Instantaneous spike frequencies of LP neurons as membrane potential is ramped before and after applying 3×10^−7^MCCAP at 10°C (left panel) and 20°C (right panel) from cold-acclimated animals(a) and hot-acclimated animals (b). c. Population averages of instantaneous spike frequencies at 10°C averaged over the −45 to −50mV range. d. Population averages of instantaneous spike frequencies at 20°C averaged over the −35 to −40mV range. Individual circles connected by lines are single animals. Horizontal bars represent population means. * p<0.05, ** p<0.01, *** p<0.001

At 20°C, 2/4 of the preparations from cold-acclimated animals and 1/5 preparations from hot-acclimated animals failed to produce spikes with current ramps (Fig 4a, b; right panels-flat lines). In the presence of CCAP however, all preparations from both groups produced spikes with the same ramps; and spike frequencies increased across the membrane potentials tested. Spike frequencies were quantified in the −40 to −30mV window; and did not differ significantly between acclimation groups at 20°C (p = 0.18) (Fig 4d). CCAP significantly increased spike frequency in LP neurons from cold-acclimated animals (Δ=15.3Hz, p = 0.03) and from hot-acclimated animals (Δ=35.4Hz, p=0.00001). However, the spike frequencies were higher in the hot acclimation group in CCAP compared to the cold acclimation group (p= 0.003).

### Computational models suggest role for modulator receptor expression in excitability differences due to acclimation

What biophysical mechanisms might account for the pronounced changes in threshold and excitability seen in response to acclimation and CCAP application? To help answer this question, we constructed semi-realistic conductance-based models of STG neurons. The effects of adding an I_MI_ current to different compartments were compared.

### Spike thresholds are temperature insensitive in a single compartment computational model

A temperature sensitive, single-compartment model of the LP cell, loosely based on previous models [50] [51] was stimulated with a current injection from 0 to 65nA over 5000ms of model time at different temperatures. The spike threshold was insensitive to either temperature or addition of I_MI_ at either temperature (Fig 5a-d).

**Fig. 5:**
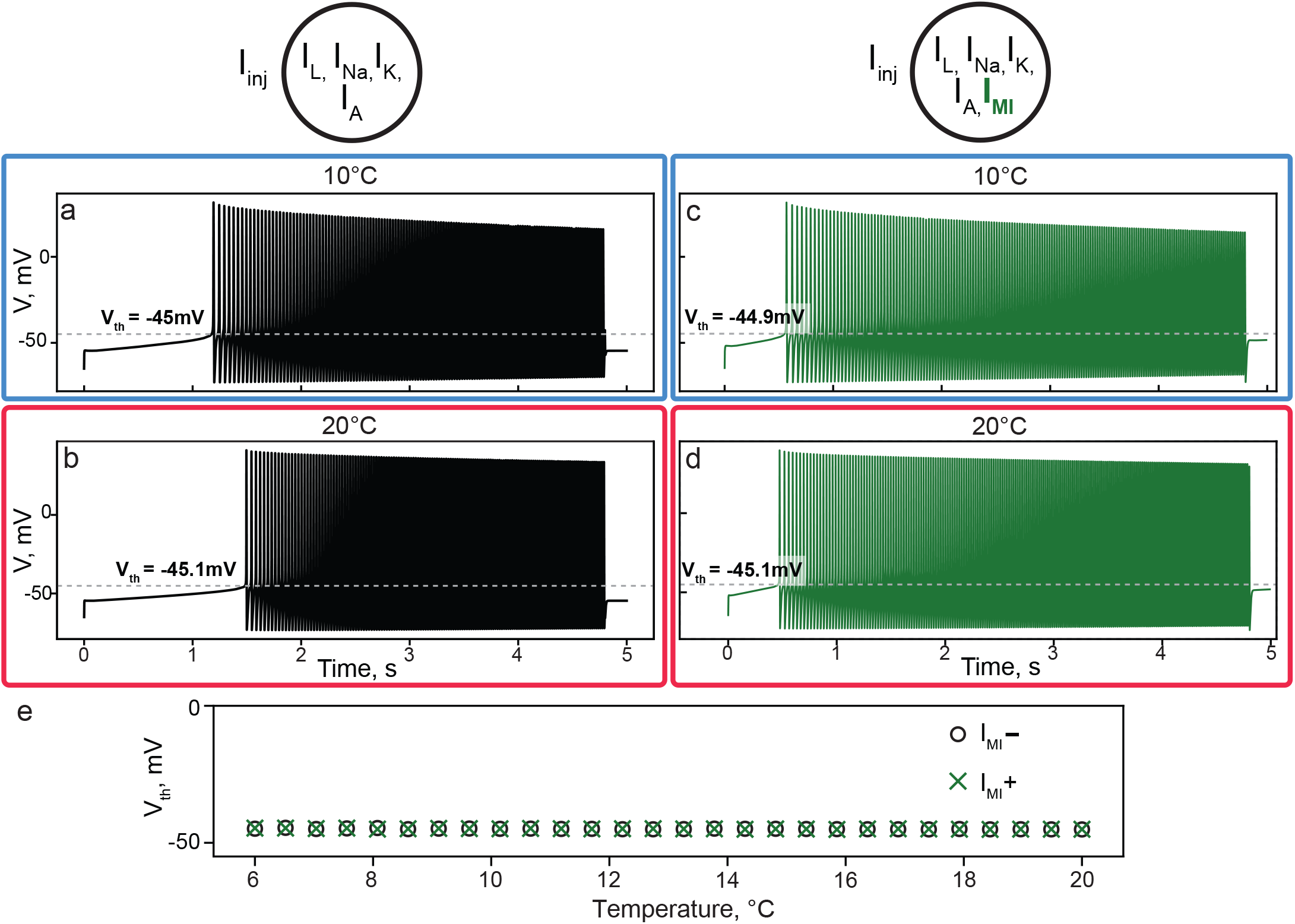
Evolution of firing threshold in a single compartment model. a, b: Voltage traces and firing thresholds of a single compartment model without I_MI_ at 10°C(a) and 20°C(b). c, d: Voltage traces and firing thresholds of a single compartment model with I_MI_ at 10°C(c) and 20°C(d). Dashed lines indicate voltage threshold of spiking. e: Evolution of firing threshold with respect to temperature for the I_MI_+ and I_MI_-models.

What differed in the various conditions is the amount of injected current needed to reach threshold because of a temperature dependent increase in leak current, corresponding to a decrease in input resistance. Without I_MI_, the models showed a distinct increase in the amount of current needed to elicit a spike from 10°C to 20°C which is reflected in the time at which the first spike was elicited (it was ramped at the same rate in all conditions) (Fig. 5a, c).

Table 2 shows that selectively removing the temperature dependence from the leak current essentially eliminated the shift in current threshold in the single compartment model. In the presence of I_MI_, the single-compartment model showed little or no change in the current threshold from 10°C to 20°C (Fig. 5b, d). The voltage threshold of spiking remained essentially constant across temperatures in both the I_MI_ and zero I_MI_ cases (Fig. 5e). I_MI_ can compensate for the temperature-dependent increases in leakiness, but the single compartment model failed to fully capture the biological observations.

**Table 2:**
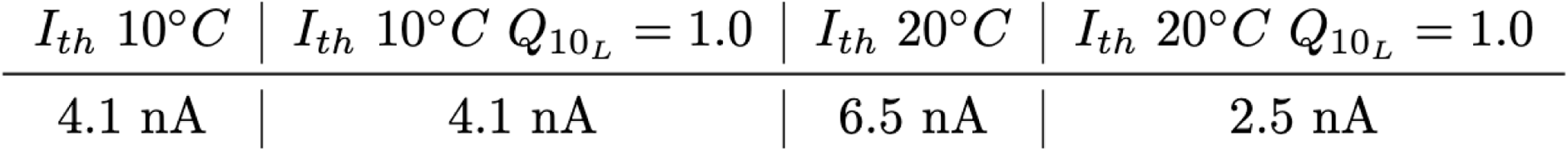
Injected current to reach spike thresholds at different temperatures.

### Temperature drives depolarization of spike thresholds in a two-compartment

#### model

Although it is often sufficient to construct single-compartment models of STG neurons to capture many of the features of their behavior [52–54], STG neuron action potentials are generated where the axons leave the neuropil and not in the somata [55]. Multi-compartmental models capture features of STG neuron function [50, 51] that separate the spike-initiating compartment from the soma compartment. We implemented a two-compartment model with a soma and an axon, which segregated the spiking generating mechanism to the axon.

We calculated the spike thresholds that would be measured in the soma and axon when current is injected into the somatic compartment at 10°C and 20°C, and then again at both temperatures with the addition of I_MI_ in the axonal compartment. The axonal compartment operates like the single compartment model. The axonal spike threshold is barely altered in the absence of I_MI_, going from 10°C to 20°C, even when the current threshold increased (Fig. 6a, b). By Ohm’s law, a more depolarized somatic potential is needed to increase a change in temperature the current reaching the axon from the soma. Consequently, the soma spiking threshold depolarized from −24.9 mV to −17.4 mV from 10°C to 20°C (Fig.6 a,b). With axonal I_MI_, this depolarization of soma spiking threshold and the current threshold increase in the axon were abolished (Fig.6c,d). Synaptic inputs that would converge to drive influence are also compartmentalized in STG neurons [56–59]. Therefore, the separation of soma from the spike initiation zone is a likely reason for the temperature driven reduction of excitability in LP neurons.

**Fig. 6:**
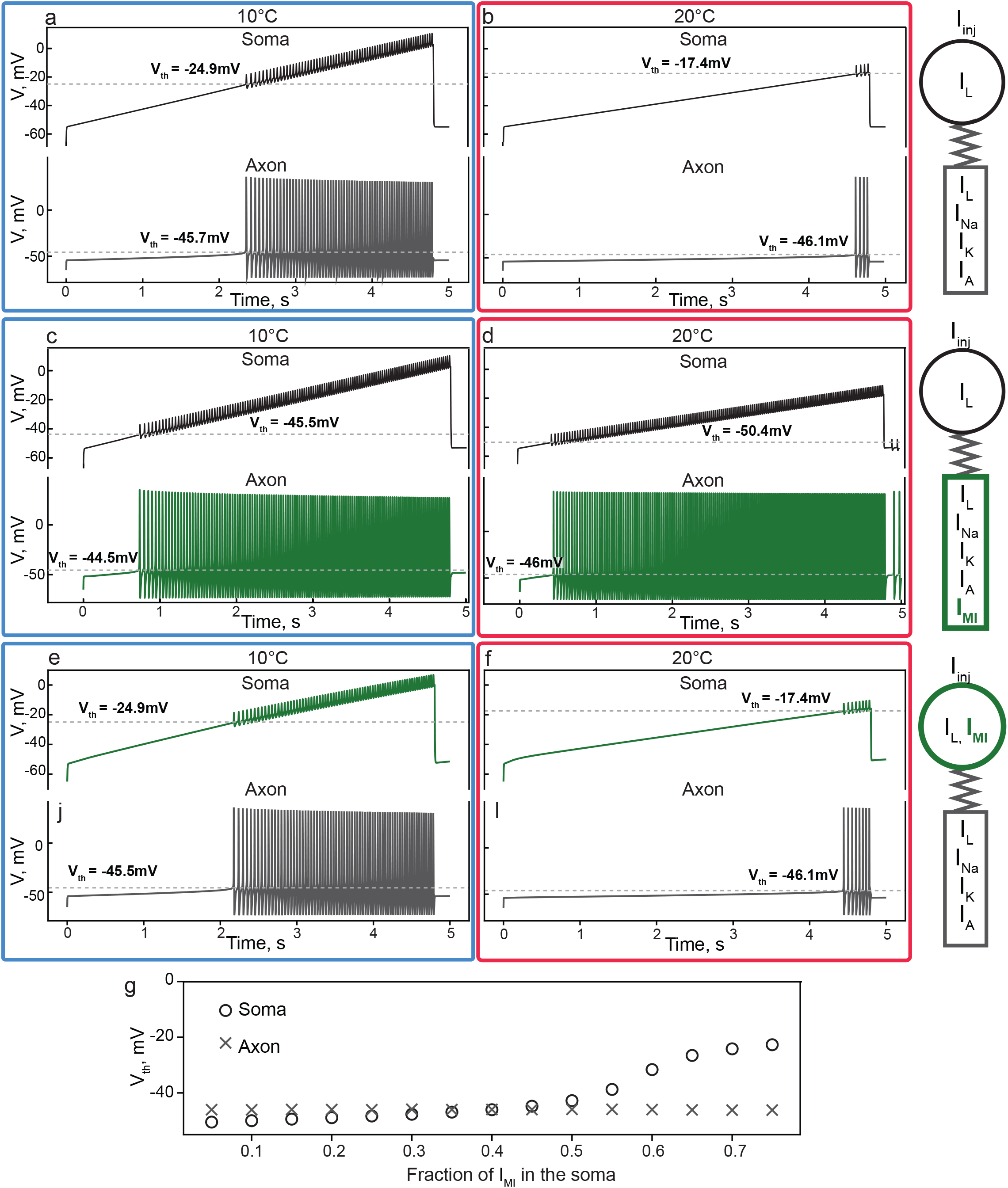
Voltage traces and firing thresholds of two-compartment models. a: No I_MI_ model at 10°C, axon (top) and soma (bottom); b: and No I_MI_ model at 20°C. c. Axonal I_MI_ model at 10°C axon (top) and soma (bottom) d. Axonal I_MI_ model at 20°C e. Soma I_MI_ model at 10°C axon (top) and soma (bottom).f. Soma I_MI_ model at 20°C. Dashed grey lines indicate voltage threshold of spiking. g: Firing thresholds at 20°C at the axon and soma in models with varying I_MI_ localization. A given fraction X of soma I_MI_ corresponds to (1-X) of I_MI_ conductance on the axon.

### Axonal distribution of I_MI_ is crucial for recovery of spike thresholds

We then examined the role of the spatial distribution of I_MI_ in the two compartments in modulating the spiking threshold. We held the two-compartment model at 20°C and compared the spiking behavior in several configurations. The models with no I_MI_ (Fig. 6b) and with I_MI_ only in the soma (Fig. 6e) showed identical voltage spiking thresholds of −17.4 mV. On the other hand, the model with I_MI_ on the axon (Fig. 6d) exhibited a more hyperpolarized spiking threshold of −50.4 mV. This suggests that I_MI_ in close proximity to the spike initiation zone enables the rescue of the spike threshold measured experimentally. We then allowed the fraction of I_MI_ in the soma vs. the axon to vary over a wider range, while recording the soma and axon thresholds for each I_MI_ distribution at 20°C (Fig. 6g). We found that spike threshold was relatively insensitive to changes in I_MI_ distribution until the fraction of axonal I_MI_ dropped to 50%. For fractions of soma I_MI_ greater or equal to 50%, the firing threshold of the soma smoothly rose towards −17.4 mV. These theoretical results further suggest that I_MI_ amplitude in the soma plays at best a limited role in determining the spike threshold.

### I_MI_ amplitude regulates spike frequencies

The experimental results suggest that the amplitude of I_MI_ was larger in neurons from the hot-acclimated group (Fig. 2).

To explore this further, we tested the effect of changing the maximal conductance of I_MI_ in the two-compartment model at 10°C and 20°C. Consistent with analysis of the model presented above, which shows the minimal impact of I_MI_ in the soma (Fig. 6), we restricted our analysis of frequency to the case where I_MI_ is expressed solely on the axon with varying maximal conductance.

The model was run with no I_MI_ as a control, as well as with I_MI_ at the full maximal conductance used throughout the rest of this study, and with I_MI_ scaled to 50 and 75%. These values were chosen because they span the region previously shown (Fig. 6m) to have essentially constant spike thresholds at the soma (75% and 100%), along with the edge of the region where spike threshold begins to depolarize (50%). This was done at both 10°C and 20°C. The spike times were identified, along with a baseline voltage trace, and an instantaneous spike frequency calculated for each voltage value.

The results closely follow the experimental evidence. At both temperatures, all three I_MI_+ conditions have higher spike frequencies at every voltage than the control, I_MI_-condition. Furthermore, the higher conductance models show upwards shifts in their frequency-voltage curves, which mirrors a similar shift seen between cells recorded from cold and hot-acclimated animals in Fig. 4c. This shift existed at 10°C, but was much larger at 20°C.

**Fig. 7:**
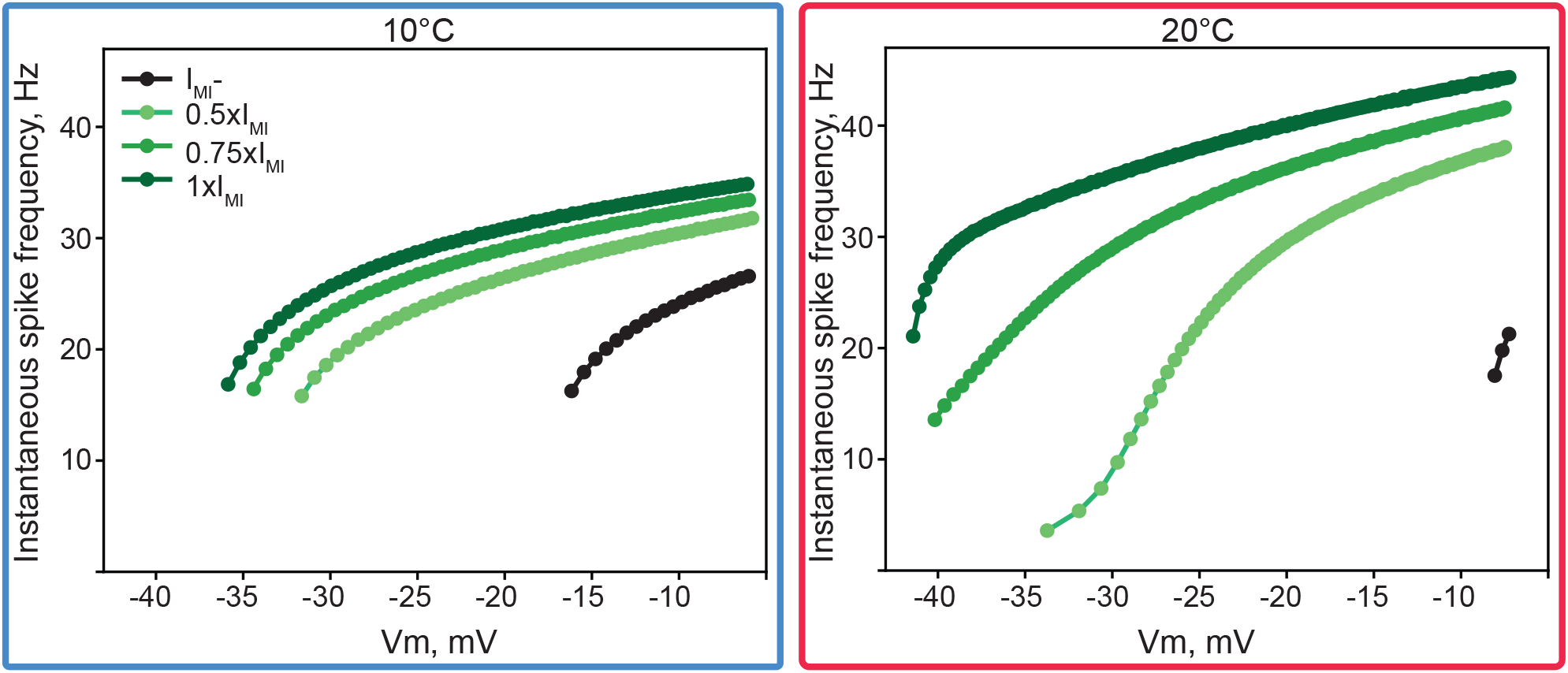
Spike frequency depends on temperature and I_MI_ amplitude in a two-compartment model. Frequency-Voltage curves for four I_MI_ conditions at 10°C and 20°C. a: f-V curves for the model at 10℃. b: f-V curves for the model 20°C Each curve represents the frequency response of the model at different baseline voltages. Green traces represent the model run at increments of I_MI_ maximal conductance, while the black trace represents the model without I_MI_. In each case, the leftward beginning of the f-V curve corresponds to the spike threshold voltage for that I_MI_ condition and temperature.

## Discussion

Stomatogastric ganglion circuitry has been studied in many decapods showing malleability of its thermal tolerance following acclimation [27, 28, 30]. Nonetheless, the processes by which the network achieves are not clear. We suggest that the neuronal mechanisms underlying acclimation should be seen as an independent form of plasticity, robustness tuning, such that its distinct triggers, mechanisms and outcomes can be investigated. Resilience to altered temperature environments likely arises from multi-dimensional interactions that have effects on numerous processes, making them difficult to untangle.

### Acclimation occurs in the context of neuronal and circuit degeneracy

It is now clear from both computational and experimental studies that similar network performance can result from multiple sets of the parameters that determine all aspects of neuronal and circuit performance [60–64]. It has also been long known that descending neuromodulatory inputs maintain rhythmic activity in the stomatogastric ganglion, and that removal of these inputs results in substantial reductions in network performance [65–68]. This suggests that decentralization and loss of activity moves preparations from the regions of parameter space in which rhythmic activity is possible, to other regions of parameter space in which are inconsistent with rhythmic activity [52]. One of the fascinating findings of the present study is that decentralization of the cold acclimated animals results in their loss of activity, but that hot acclimated animals retain rhythmic activity when decentralized. This suggests that the hot acclimation animals have retuned their sets of synaptic and intrinsic conductances and found their way into the space of degenerate solutions consistent with pyloric activity.

Interestingly, retuning of parameters to recover a desired activity profile is thought to account for recovery of activity in the pyloric rhythm after long-term decentralization [68–71], although this retuning has been attributed to homeostatic mechanisms, which we argue, may differ substantively from those triggered by acclimation.

### Acclimation impacts acute temperature sensitivity

Previous work with the crab, *Cancer borealis*, had demonstrated that both acclimation in the ocean [72] and in laboratory tanks [28] altered the robustness of the pyloric rhythm to acute temperature changes. In these studies, the range over which pyloric rhythms were found was extended by warm acclimation. The results in this paper show similar effects for *H. americanus*. Interestingly, in both species, the upper range of robust pyloric rhythms was extended by at least 7-12°C, and at their upper ranges, the pyloric rhythms were faster than had been previously reported, although they retained an appropriate triphasic rhythm. As maintaining a triphasic rhythm over a large frequency range likely involves a number of cellular and synaptic mechanisms [20, 73], it is likely that many features of the pyloric neurons and synapses must be coordinately regulated to produce these high frequency triphasic rhythms.

### The interaction between neuromodulation and acclimation

The STG is modulated by a large number of peptides and amines [36, 74], among them CCAP, which we have studied here. It has been previously shown that the responses of decentralized preparations to acute temperature ramps are dramatically altered by application of exogenous modulator [75–77]. Unlike many neuromodulators that are released by descending modulatory neuron and/or sensory neurons, CCAP is hormonally released, and reaches the STG via the circulation. Thus, decentralization does not change the concentrations of CCAP that reach the STG, as removal from the stomach has already removed all sources of CCAP from the dissected preparations. Nonetheless, the data in this paper show that exogenously applied CCAP can only partially overcome the loss of rhythmic activity in cold-acclimated animals seen at higher temperatures, and CCAP-treated cold acclimated preparations do not reach the level of activity seen when CCAP is applied to warm-acclimated animals.

Preparations from cold and warm acclimated animals showed significant differences in their CCAP dose-response curves and single cell excitability in response to CCAP. It is tempting to therefore conclude that CCAP receptor number, distribution, or signal transduction pathways are directly influenced by acclimation. This interpretation is consistent with the computational model that shows that some of the acclimation effects on LP neuron threshold result from changes in I_MI_ distribution. But, as with all functional physiological dose-response curves, it is not straightforward to distinguish between changes in CCAP number and distribution from other aspects of the physiological properties of the target neurons through which we see the effects of the agonist. Therefore, it will be interesting to obtain more direct measurements of CCAP receptors, the I_MI_ changes and all of the other ion channels that contribute to the CCAP effects.

### Robustness Tuning, Learning, and Homeostasis

We suggest that acclimation should best be considered a form of robust tuning and distinguished from other forms of plasticity such as learning and homeostasis of intrinsic excitability and synaptic scaling. Acclimation is engaged in ethological circumstances to produce long-lasting changes that are reversible. The lobsters we studied here can live for more than twenty years and therefore must repeatedly adapt to substantial seasonal changes in ocean temperature[78]. Acclimation must persist over time scales of months but also must reverse over times scales of months in the wild. In contrast, in some instances, learning creates an almost permanent change in brain function that persists throughout the animal’s lifetime, and homeostatic processes are intended to be operational to maintain both short-term and long-term stability in the face of multiple perturbations. It is likely that all three forms of plasticity engage use some of the same molecular and cellular mechanisms, but it is quite mysterious how these different forms of plasticity coexist in the same neurons and circuits to maintain resilient and robust nervous system function in response to the complex sets of perturbations that animals encounter in their natural worlds.

## Methods

### Experimental Animal and Preparation Details

Adult Americanus lobsters (*Homarus americanus*) were purchased from Commercial Lobster (Boston, MA, USA) and held in artificial seawater at 4°C or 18°C on a 12 h light/12 h dark cycle for a minimum period of 3 weeks. Animals were placed on ice for 30 min prior to dissection. The stomach was dissected from the animal. The intact stomatogastric nervous system (STNS) was isolated from the stomach, including: the two bilateral commissural ganglia, esophageal ganglion, and STG, with connecting motor nerves. The STNS was pinned onto a Sylgard-coated Petri dish (10 mL) and continuously superfused with chilled saline (composition in mM: 479.12 NaCl, 12.74 KCl, 13.67 CaCl_2_, 20.00 MgSO_4_, 3.91 Na_2_SO_4_, 11.45 Trizma base, and 4.82 maleic acid; pH 7.45 at room temperature ∼23°C).

### Electrophysiology

The STG was desheathed and intracellular recordings from somata were performed with 5-20 MΩ glass microelectrodes filled with 0.4 M K_2_SO_4_, 3.5 mm NaCl, and 25 mm KCl. Intracellular signals were recorded with an Axoclamp 900A amplifier (Molecular Devices, San Jose, CA, USA). Neuronal identity was established after impaling the somata with sharp electrodes based on spiking activity observed on their respective nerves. Extracellular nerve recordings from the STG were made by building wells around nerves using a mixture of Vaseline and mineral oil and placing stainless steel pin electrodes within the wells to monitor spiking activity. Extracellular nerve recordings were amplified using model 3500 extracellular amplifiers (A-M Systems). Data were acquired using a Digidata 1440 digitizer (Axon Instruments) and pClamp data acquisition software (Axon Instruments, version 10.7). Removal of neuromodulatory inputs to the STG (decentralization), was performed by adding 0.1 μM tetrodotoxin (TTX) to a Vaseline well built around the stomatogastric nerve. Current injection ramps were performed in the presence of 10^−5^ M picrotoxin (PTX) superfused continuously. For current injections neurons were impaled with two electrodes, one of which was used for current injections and the second for reporting membrane potential.

### Pharmacology and temperature control

Temperature of superfused saline solution was controlled using an SC-20 Peltier device with a model CL-100 temperature controller (Warner Instruments). Temperature was monitored using a Warner Instruments TA-29 thermocouple placed within 1 cm of the STG. The temperature was manually increased by 1°C/min for the temperature ramps starting at 11°C. Crustacean cardioactive peptide (CCAP) was purchased from Targetmol Chemicals Inc, and Bachem (TA9H9877110F, 4015114). Stocks were made in water and diluted in saline to the appropriate concentrations for use. For dose response curves doses were applied serially with no intervening washes and repeated at two temperatures in the same preparations. Dish temperature was set to either 10 or 20°C for the first dose response in a random order. For current ramps, cells were first held at 20°C, a single dose (300nM CCAP) was applied for 10 minutes and washed out for >30 minutes before dropping the temperature to 10°C and repeating the process.

### Electrophysiological and Statistical Analyses

Electrophysiological data were analyzed with custom MATLAB scripts to calculate burst, spike properties and triphasicity. Spikes from extracellular recordings were semi-manually identified and assigned to individual neurons using Crabsort (https://github.com/sg-s/crabsort). Bursts were identified as having 3 or more spikes. Dose response properties are averaged over all bursts in the last two minutes of peptide application. Percentage of triphasic transitions was calculated using the start times of each PD, LP, and PY burst in a two-minute stretch of the recording for each dose. Each transition consisted of a pair of adjacent bursts. A transition was marked as triphasic if the transitions were any of the following: PD before LP, LP before PY, or PY before PD. The count of triphasic transitions was then divided by the total number of transitions (number of bursts – 1) to get the final percentage. To calculate spike threshold, spike onset was defined as the voltage corresponding to the maximum curvature of the first derivative of the voltage (dV/dt), when dV/dt crossed the threshold value of 10 mV/ms with manual corroboration. Spike frequencies during current ramps were computed from manually curated spike times and were not redetected. A 0.20 s moving median filter was used to suppress spikes and estimate a smooth baseline membrane-potential trace followed by Savitzky–Golay smoothing using a 0.50 s window and third-order polynomial and manually verified. Instantaneous spike frequencies were calculated as the reciprocal of the inter-spike interval and the time of the second spike was used to extract membrane potential at the frequency. Instantaneous-frequency values were a sliding average of ±2 mV bins of baseline voltage. Silent ramps, or ramps with fewer than two spikes, were plotted as 0 Hz.

### Temperature versus burst frequency statistics

Burst frequency for each animal was averaged at each temperature. Linear mixed-effects model were tested with temperature, acclimation group (hot vs cold), and their interaction as fixed effects, and animal identity as a random effect to account for repeated measures. Post-hoc comparisons between hot- and cold-acclimated animals at each temperature were performed using estimated marginal means with Holm correction for multiple comparisons.

### Dose response statistics

Effects of acclimation group, temperature, and CCAP were analyzed using a linear mixed-effects model with acclimation group, CCAP dose, and temperature as fixed effects and animal as a random effect. Following model fitting, Type III ANOVA was performed to evaluate main effects and interactions. Degrees of freedom were estimated using the Satterthwaite approximation. Planned post-hoc comparisons between hot- and cold-acclimated animals at each temperature and condition were performed using estimated marginal means with Holm corrections.

Excitability statistics: Spike threshold data were analyzed using a linear mixed-effects model. The model included acclimation condition, ramp temperature and neuromodulator state (baseline vs CCAP), as fixed effects, with animal as a random intercept to account for repeated measures. Following model fitting, a Type III ANOVA was performed to evaluate main effects and interactions. Degrees of freedom were estimated using the Satterthwaite approximation. Since a significant modulator x ramp temperature interaction was detected, post-hoc comparisons were conducted using estimated marginal means with Kenward–Roger degrees of freedom correction. Simple effects were examined to determine how drug effects varied across temperature and acclimation groups.

Instantaneous spike frequencies were analyzed separately at 10°C and 20°C. Within each temperature, baseline and CCAP conditions were compared using paired t-tests within hot- and cold-acclimated groups. Between-group comparisons (hot vs cold) were conducted using Welch’s independent-samples t-tests at baseline and during CCAP. All tests were two-tailed.

### Computational Models

Mathematical models of the LP cell were constructed to better understand the dynamics observed in the experimental results. All models were built based on the Hodgkin-Huxley formalism and loosely informed by voltage clamp data drawn from Schneider et al in *C. borealis*.

### Single Compartment Model

A family of single-compartment models was constructed. These models varied in temperature and the presence or absence of an I_MI_ current. This latter variable was used to model the effect of CCAP on decentralized preparations.

The model contained a fast sodium current (I_Na_), a delayed rectifier potassium current (I_Kd_), a transient potassium channel (I_Kd_), a leak channel (I_L_), and a modulator-dependent inward current (I_MI_) which was used to model the electrophysiological impact of CCAP in the experimental results. In addition, each simulation included an injected current ramp (I_Inj_) which ran from 0-20 nA over 5000 ms of model time.

The single-compartment models obeyed the current balance equation

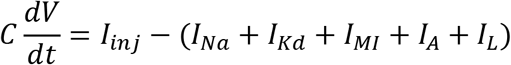

where each ionic current except I_MI_ had the standard form

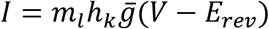

with *l* and *k* were integers. *m* and *h* obey an equation with the form and *m*_∞_, *h*_∞_ are expressed as variations (given in Table 1) on a sigmoid with the standard form

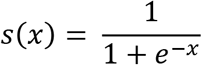

and *τ*(V) is a voltage dependent time scale, defined for each current in Table 1. Conductance parameters

| Currents | Gates | $m_\infty$ | $\tau_m$(ms) | $h_\infty$ | $\tau_h$(ms) | $\bar{g}$(nS) | $E_{rev}$(mV) |
| --- | --- | --- | --- | --- | --- | --- | --- |
| $I_A$ | $m^2 h$ | $s(\frac{v+54}{10})$ | 15 | $s(-\frac{v+60}{5})$ | 100 | 0.059 | -80 |
| $I_{Kd}$ | $m^4$ | $s(\frac{v+23}{5})$ | $2 + 14s(-\frac{v+23}{5})$ | - | - | 17.9 | -80 |
| $I_{Na}$ | $m^3 h$ | $s(\frac{v+18}{12.25})$ | 2 | $s(-\frac{v+28}{7.7})$ | 5 | 80 | 50 |
| $I_{MI-P}$ | $m$ | $s(\frac{v+19}{18})$ | 2 | - | - | 0.283 | -10 |
| $I_{MI-T}$ | $m^3 h$ | $s(\frac{v+52.5}{4.8})$ | $2 + \frac{150}{\cosh(\frac{v+56}{1.6})}$ | $[s(-\frac{v+61}{2.8}) - 0.1]^+$ | $100 + \frac{600}{\cosh(0.25(v+60))} + 200s(0.5(v+60))$ | 2.83 | -16 |
| $I_L$ | - | - | - | - | - | 0.795 | -55 |

Unlike the other current in the single compartment model, the I_MI_ current is split into a persistent and a transient component. Each of these components, described in Table 1, individually follows the form given above. The total I_MI_ current is then given as

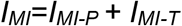

### Two Compartment Model

The two-compartment model followed the current balance equations

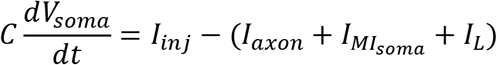

And

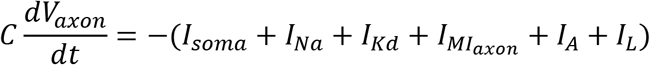

Where *I*_axon_ and *I*_soma_ are the axial conductances connecting the two compartments, with the forms

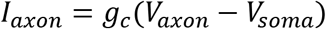

And

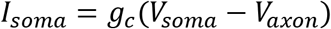

respectively.

The currents had the same forms as those in the single-compartment model, given in Table 1, with the exception of I_MI_, which is divided into two currents, one for each compartment.

In addition, each simulation included an injected current ramp (I_inj_) which ran from 0-65 nA over 5000 ms of model time.

### Temperature Dependence

In both the single, and two-compartment models, temperature dependence was expressed by applying Q_10_ factors to the maximal conductances and the current time scales as follows.

Maximal conductances scaled with temperature according to the following rule

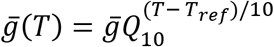

While time scales were scaled in the following way

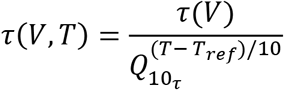

Where T_ref_ was 10°C, Q_10g_ was 1.6, and Q_10_*_ττ_* was 3.0, and in the two-compartment model the temperature dependence of the axial conductance Q_10c_ was 1.15. These values were chosen based on values given in previous studies.

### I_MI_

The two compartment model was constructed with I_MI_ in both the axon and soma. The activation and inactivation characteristics of these currents were modeled identically but independently, with each compartmental I_MI_ current depending on its voltage of its local compartment.

In addition, the model allowed us to vary the maximal conductance of the I_MI_ on each compartment. This was done by assigning a fixed percentage of the *g* to each compartment, such that together they summed to the full *g_MMM_*.

### Computational Details

All models were implemented as custom python code. Simulations were solved using Scipy’s solve_ivp function, running for 5000 ms of model time.

---

**Supplementary figure S1.**
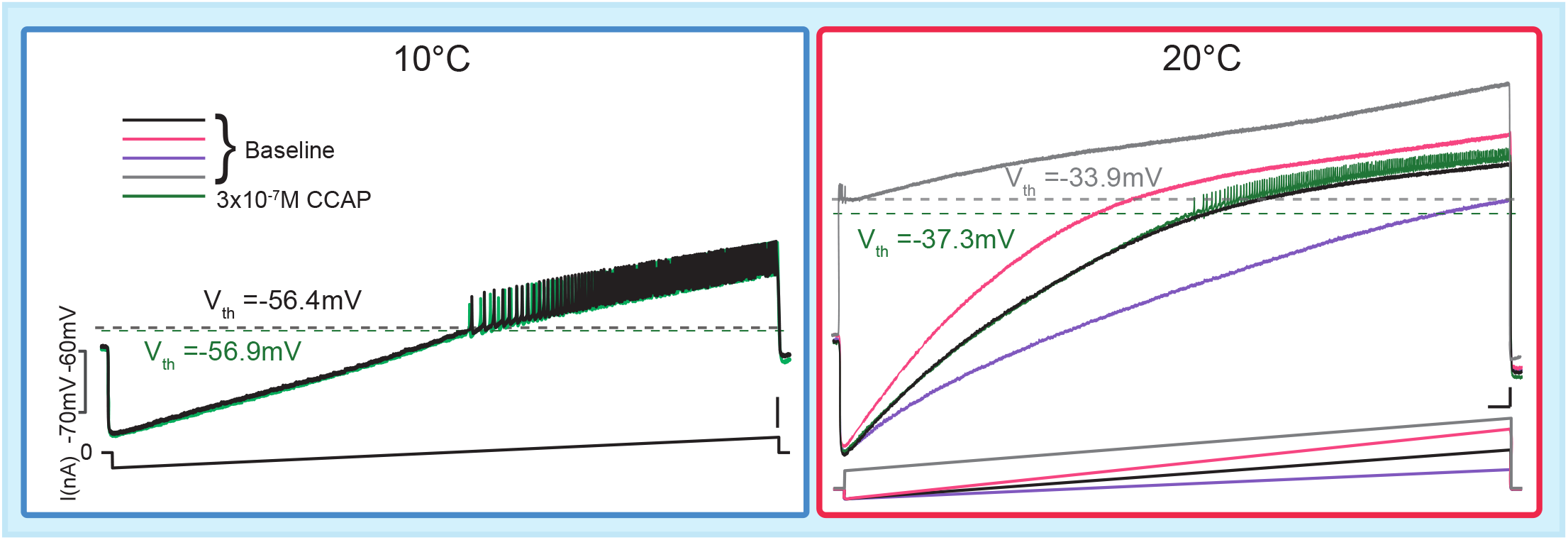
Spike failure with ramps at high temperatures: Sample traces from a cold animal of membrane voltage (top traces) and current injections (bottom traces). Both voltage traces are to the same current injection in the left panel. Right panel the colors of the voltage traces match current traces they’re paired with. CCAP was applied with the black current injection.

**Supplementary table S1.** Mean±S.D. values for PD burst frequency in cold and hot acclimated animals. Significant effect of temperature on firing frequency, p < 0.0001, from 11°C to 23°C.

| Temp (°C) | 11 | 12 | 13 | 14 | 15 | 16 | 17 | 18 | 19 | 20 | 21 | 22 | 23 |
| --- | --- | --- | --- | --- | --- | --- | --- | --- | --- | --- | --- | --- | --- |
| Cold | 0.57 ± 0.17 | 0.64 ± 0.16 | 0.7 ± 0.18 | 0.8 ± 0.21 | 0.87 ± 0.23 | 0.94 ± 0.26 | 0.92 ± 0.3 | 1.03 ± 0.33 | 1.09 ± 0.35 | 1.18 ± 0.37 | 1.15 ± 0.36 | 1.19 ± 0.36 | 1.22 ± 0.36 |
| Hot | 0.65 ± 0.09 | 0.72 ± 0.11 | 0.84 ± 0.09 | 0.93 ± 0.09 | 1.01 ± 0.09 | 1.09 ± 0.13 | 1.2 ± 0.15 | 1.3 ± 0.17 | 1.4 ± 0.21 | 1.4 ± 0.23 | 1.5 ± 0.26 | 1.6 ± 0.27 | 1.6 ± 0.3 |

**Supplementary table S2.** Pairwise Comparisons (Cold acclimated–Hot acclimated at each Temperature): Linear mixed effects model run. No main effect of acclimation group, significant acclimation x temperature interaction (F(4,94.265) = 4.3357, p < 0.0001). Post-hoc comparisons performed with Holm correction.

| Temp (°C) | 11 | 12 | 13 | 14 | 15 | 16 | 17 | 18 | 19 | 20 | 21 | 22 | 23 |
| --- | --- | --- | --- | --- | --- | --- | --- | --- | --- | --- | --- | --- | --- |
| p | 0.46 | 0.32 | 0.17 | 0.1 | 0.1 | 0.022 | 0.006 | 0.002 | 0.002 | 7E-04 | 1E-04 | <0.0001 | <0.0001 |

**Supplementary table S3.** Mean±S.D. values for triphasicity %.

| Temp | Dose<br>Acclim. | Baseline | 3x10<sup>-9</sup><br>CCAP | 10<sup>-8</sup><br>CCAP | 3x10<sup>-8</sup><br>CCAP | 10<sup>-7</sup><br>CCAP | 3x10<sup>-7</sup><br>CCAP | 10<sup>-6</sup><br>CCAP |
| --- | --- | --- | --- | --- | --- | --- | --- | --- |
| 10°C | Cold | 0.263 ± 0.163 | 0.230 ± 0.231 | 0.391 ± 0.263 | 0.827 ± 0.262 | 0.989 ± 0.199 | 0.913 ± 0.179 | 0.940 ± 0.142 |
|  | Hot | 0.609 ± 0.316 | 0.856 ± 0.283 | 0.998 ± 0.005 | 1 ± 0 | 1 ± 0 | 1 ± 0 | 1 ± 0 |
| 20°C | Cold | 0 ± 0 | 0 ± 0 | 0 ± 0 | 0.240 ± 0.374 | 0.531 ± 0.405 | 0.520 ± 0.382 | 0.591 ± 0.435 |
|  | Hot | 0.120 ± 0.173 | 0.176 ± 0.183 | 0.292 ± 0.269 | 0.821 ± 0.102 | 0.996 ± 0.009 | 1 ± 0 | 1 ± 0 |

**Supplementary table S4.**
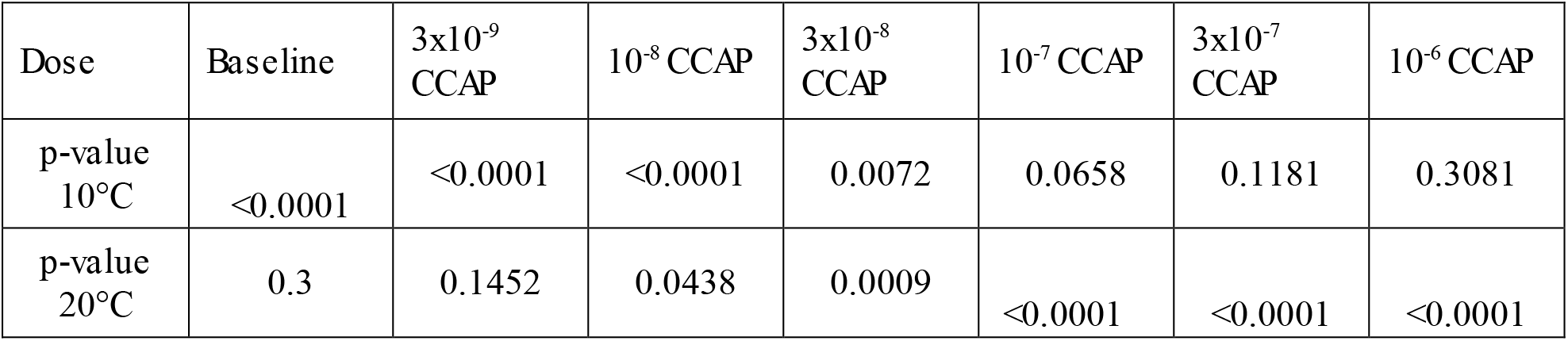
Pairwise comparisons of triphasicity % between cold and hot acclimated groups: Linear mixed effects models present significant main effects of acclimation group (F(1,10) = 15.48, p = 0.0028), CCAP dose (F(1,30) = 85.88, p < 0.0001), and temperature (F(1,30) = 35.95, p < 0.0001. A significant acclimation x CCAP dose x temperature interaction (F(1,30) = 14.2253, p = 0.0007123) emerged. Adjusted p = 0.0004, for groupwise comparisons at 10°C. Adjusted p < 0.0001 for groupwise comparison at 20°C. Post-hoc comparisons with Holm correction.

**Supplementary table S5.**
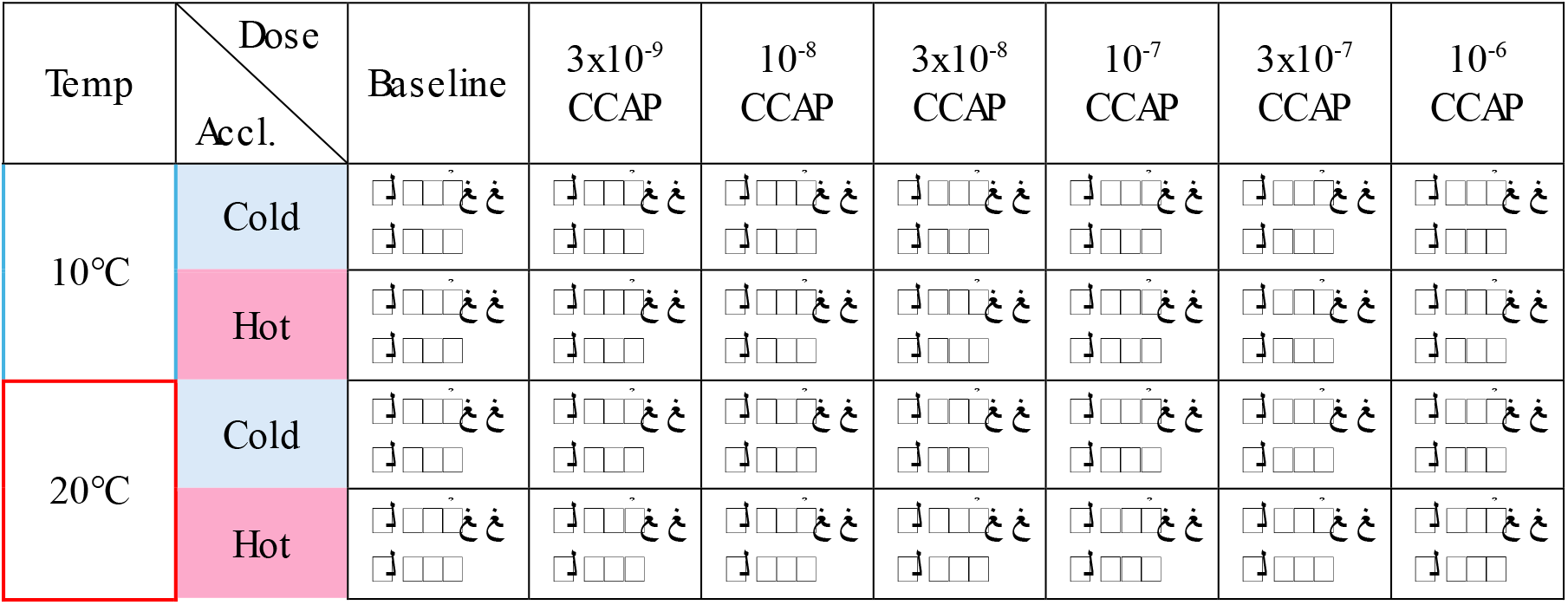
Mean±S.D. PD burst frequency values.

**Supplementary table S6.**
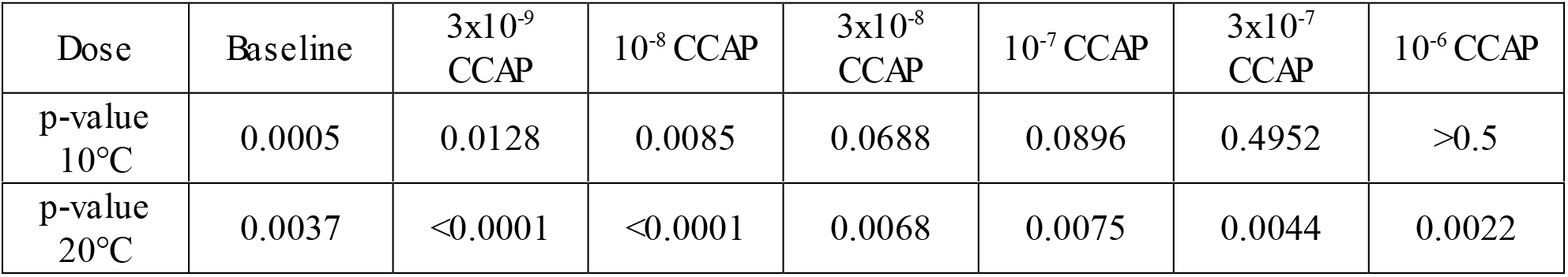
Pairwise comparisons between cold and hot acclimated animal PD burst frequency values: 10°C-There was a significant main effect of acclimation group (*F*(1,12) = 9.62, *p* = 0.0092) (PD burst frequencies are different between hot and cold) and CCAP dose (*F*(5,60) = 2.49, *p* = 0.041).CCAP dose impacted PD burst frequency. There was a significant interaction between acclimation group and CCAP dose (*F*(5,60) = 3.25, *p* = 0.0115). 20°C-There was a significant main effect of acclimation (F(1,12) = 27.84, p = 0.0002) and CCAP dose (F(6,72) = 9.46, p < 0.0001. No significant interaction between dose and acclimation, there were similar CCAP-driven increases in both groups.

**Supplementary table S7.** Mean±S.D. values for LP spike frequency.

| Temp | Dose<br>Accl. | Baseline | 3x10<sup>-9</sup><br>CCAP | 10<sup>-8</sup><br>CCAP | 3x10<sup>-8</sup><br>CCAP | 10<sup>-7</sup><br>CCAP | 3x10<sup>-7</sup><br>CCAP | 10<sup>-6</sup><br>CCAP |
| --- | --- | --- | --- | --- | --- | --- | --- | --- |
| 10°C | Cold | 0±0 | 0.112±<br>0.161 | 1.146±<br>1.791 | 3.687±<br>2.229 | 4.065±<br>2.494 | 5.022±<br>3.009 | 5.651±<br>3.1 |
|  | Hot | 1.311±<br>1.159 | 1.786±<br>1.049 | 2.607±<br>0.718 | 4.405±<br>1.001 | 5.961±<br>0.964 | 6.908±<br>1.132 | 6.381±<br>1.827 |
| 20°C | Cold | 0±0 | 0±0 | 0±0 | 1.688±<br>2.882 | 4.468±<br>6.077 | 4.767±<br>4.652 | 6.451±<br>3.720 |
|  | Hot | 0±0. | 0±0 | 0.448±<br>0.984 | 4.158±<br>2.699 | 10.543<br>±3.676 | 12.492<br>±3.900 | 12.993<br>±3.568 |

**Supplementary table S8.** Pairwise comparisons between hot and cold animals LP spike frequencies. 10°C-There was a significant main effect of acclimation group (*F*(1,12) = 9.89, *p* = 0.0085) and CCAP dose (*F*(5,60) = 22.26, *p* < 0.0001). There was a significant interaction between acclimation group and CCAP dose (*F*(5,60) = 2.74, *p* = 0.027). 20°C-There was a significant main effect of acclimation (*F*(1,12) = 9.65, *p* = 0.0091), and CCAP dose (*F*(6,72) = 38.66, *p* < 0.0001). There was a significant interaction between acclimation group and CCAP dose (*F*(6,72) = 4.93, *p* = 0.00028).

| Dose | Baseline | 3x10<sup>-9</sup><br>CCAP | 10<sup>-8</sup> CCAP | 3x10<sup>-8</sup><br>CCAP | 10<sup>-7</sup> CCAP | 3x10<sup>-7</sup><br>CCAP | 10<sup>-6</sup> CCAP |
| --- | --- | --- | --- | --- | --- | --- | --- |
| p-value<br>10°C | 0.0014 | 0.0064 | 0.007 | 0.288 | 0.276 | 0.726 | >0.5 |
| p-value<br>20°C | 1 | 1 | 0.269 | 0.0017 | 0.0026 | 0.0027 | 0.0026 |

**Supplementary table 7.**
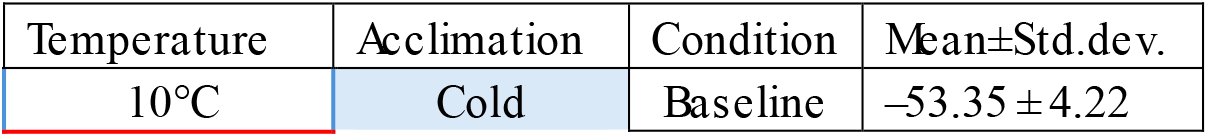
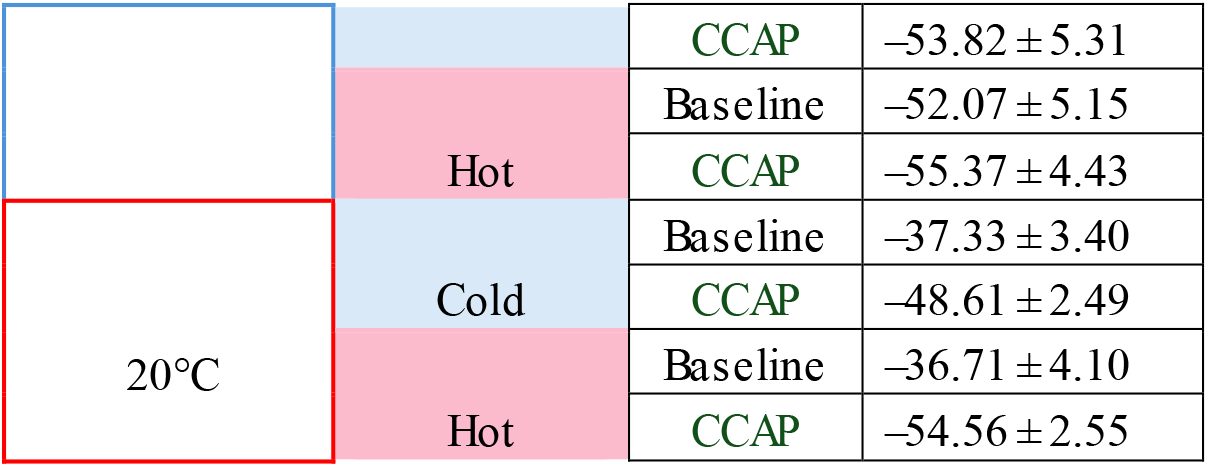
Spike thresholds, mV. The linear mixed-effects model revealed significant main effects of modulator (F(1,21)=26.18, p=4.5×10⁻⁵) and ramp temperature (F(1,21)=33.12, p=1.0×10⁻⁵). The modulator x ramp temperature interaction was also significant (F(1,21)=14.02, p=0.0012). There was no significant main effect of acclimation group (hot vs cold; p=0.31), nor any significant interactions involving acclimation (all p > 0.18).

**Supplementary table 8.** Post-hoc comparisons of spike thresholds in Baseline versus CCAP using estimated marginal means Post-hoc comparisons of baseline spike thresholds at 10°C versus at 20°C. Direct interaction contrast confirmed that the drug effect differed between 10 and 20 °C (p=0.0012).

| Temp | Acclimation | p-value |
| --- | --- | --- |
| 10°C | Cold | 0.883 |
|  | Hot | 0.211 |
| 20°C | Cold | 0.0017 |
|  | Hot | <0.0001 |

| Acclimation | p-value |
| --- | --- |
| Cold | <0.0001 |
| Hot | 0.0003 |

**Supplementary table 9.** Instantaneous spike frequencies.

| Temperature | Acclimation | Condition | Mean±Std.dev. |
| --- | --- | --- | --- |
| 10°C<br>(-45 to -50mv) | Cold | Baseline | 7.12 ± 4 |
|  |  | CCAP | 9.45 ± 3.21 |
|  | Hot | Baseline | 7.23 ± 3.03 |
|  |  | CCAP | 15.42 ± 3.94 |
| 20°C<br>(-35 to -40mv) | Cold | Baseline | 2.09 ± 2.45 |
|  |  | CCAP | 17.42 ± 7.67 |
|  | Hot | Baseline | 5.08 ± 3.6 |
|  |  | CCAP | 40.50 ± 5.8 |

**Supplementary table 10.** Paired T-test results for baseline versus CCAP comparisons of spike frequencies.

| Temp | Acclimation | p-value |
| --- | --- | --- |
| 10°C | Cold | 0.0368 |
|  | Hot | 0.0119 |
| 20°C | Cold | 0.0281 |
|  | Hot | 0.000014 |

